# Fine-mapping and identification of candidate causal genes for tail length in the Merinolandschaf breed

**DOI:** 10.1101/2022.02.27.481613

**Authors:** Dominik Karl Lagler, Elisabeth Hannemann, Kim Eck, Jürgen Klawatsch, Doris Seichter, Ingolf Russ, Christian Mendel, Gesine Lühken, Stefan Krebs, Helmut Blum, Maulik Upadhyay, Ivica Međugorac

**Affiliations:** Population Genomics Group, Department of Veterinary Sciences, LMU Munich, Lena-Christ-Str. 48, 82152 Martinsried, Germany; Tierzuchtforschung e.V. München, Senator-Gerauer-Str. 23, 85586 Poing, Germany; Institute for Animal Breeding, Bavarian State Research Center for Agriculture, Prof.-Dürrwaechter-Platz 1, 85586 Poing, Germany; Department of Animal Breeding and Genetics, JLU Gießen, Ludwigstr. 21a, 35390 Gießen, Germany; Laboratory for Functional Genome Analysis, Gene Center, Ludwig-Maximilians-University Munich, 80539 Munich, Germany

**Keywords:** sheep, lambs, tail length, docking, animal welfare, mapping study, *HOXB13*

## Abstract

**Background:** Docking the tails of young lambs in long-tailed sheep breeds is a common practice worldwide. This practice is associated with pain, suffering and damage to the affected animals. Breeding for a shorter tail in long-tailed sheep breeds could offer one of the alternatives. This study aimed to analyze the natural tail length variation in the most common German Merino variety, and to identify possible causal alleles for the short tail phenotype segregating within a typical long-tailed breed.

**Results:** Haplotype-based mapping in 362 genotyped (Illumina OvineSNP50) and phenotyped Merinolandschaf lambs resulted in a genome-wide significant mapping at position 37,111,462 bp on sheep chromosome 11 and on chromosome 2 at position 94,538,115 bp (Oar_v4.0). Targeted capture sequencing of these regions in 48 selected sheep and comparative analyses of WGS data of various long and short-tailed sheep breeds as well as wild sheep subspecies identified a SNP and a SINE element as the promising candidates. The PCR genotyping of these candidates revealed complete linkage of both the candidate variants. The SINE element is located in the promotor region of *HOXB13*, while the SNP was located in the first exon of *HOXB13* and predicted to result in a nonsynonymous mutation.

**Conclusions:** Our approach successfully identified *HOXB13* as candidate genes and the likely causal variants for tail length segregating within a typical long-tailed Merino breed. This would enable more precise breeding towards shorter tails, improve animal welfare by amplification of ancestral alleles and contribute to a better understanding of differential embryonic development.

## Background

Sedentary human communities began sheep management as early as 10,000-11,000 BP in an area stretching from central Anatolia to northwestern Iran ^1^. It is proposed that the Asiatic mouflon (*Ovis orientalis*), which was common in the same area, was the wild ancestor. The Asiatic mouflon, like other wild sheep subspecies, is a short-tailed hair sheep. Accordingly, the first domesticated sheep were also short-tailed hair sheep, kept mainly for their meat ^2^. The systematic production and processing of wool did not occur until several millennia later, leading to the “Secondary Product Revolution” ^3^ and the worldwide spread of wool sheep. Long-term selection for fine wool fibers culminated in the economically most important and widespread sheep breed, the Merino. All Merinos are characterized by long tails and the common occurrence of fine wool and long tail led to the frequent opinion that these phenotypes are also genetically coupled or the result of the same artificial selector ^4^. This assumption could not be proven directly, however, in today’s sheep husbandry systems the long woolly tail comes with several problems, e.g. the accumulation of dags in the tail area, which predisposes for flystrike ^5^. Therefore, most lambs of long-tailed breeds worldwide are docked shortly after birth ^4^. With increasing importance of animal welfare in our society and subsequent restrictions and prohibitions of practices that cause pain and suffering to animals, tail docking has come under scrutiny. In Scandinavia, tail docking without a veterinary indication has already been made illegal ^6^ and in the Netherlands, exclusions from the docking ban have only been granted for three long-tailed English breeds under the condition of an effective breeding program for shorter tails ^7^. In Austria, tail docking in lambs is allowed until an age of 7 days, provided the operation is done by a veterinarian or another qualified person and an analgesic for intra- and post-operative pain-relief is given ^8^. The German Animal Welfare Act currently still allows tail docking without anesthesia for lambs under eight days of age, but future amendments will probably seek to eliminate exceptions to the amputation ban ^9^.

These developments clearly show that a long-term non-invasive alternative for the painful practice of tail docking is urgently needed. Here, a genetic solution offers itself. A high ethical acceptance of genetic breeding for a shorter tail could be expected as all wild sheep subspecies and thus also the ancestor of today’s domestic sheep have naturally short tails ^10,11^. Therefore, this breeding could be seen as a “back to the roots” program.

The genetic basis for shorter tail breeding efforts is provided by the medium to high heritability of tail length in different sheep breeds, e.g. 0.58 in Merinos ^12^ or 0.77 in Finnish landrace sires ^13^. James, et al. ^14^ suggest that the inheritance of tail length in Australian Merino depends on a small number of interacting genes of large effects, in which short tail genes show a possible dominance. The presence of various short-tailed Nordic breeds offers a possibility of genetic reduction of tail length by introgression of the desired genetic variants from short-tailed breeds. Scobie and O’Connell ^15^ crossed short-tailed Finnsheep with long-tailed Cheviot sheep, and observed that an increased proportion of Finnsheep genes led to a proportional reduction in tail length. However, this option is unpopular with breeders, as crossing is associated with the loss of breed-specific traits and a possible decline in previously achieved breeding progress in important production traits.

As an alternative, breeders in Australia and New Zealand attempted genetic shortening of the tail by phenotypic selection within individual breeds. However, Carter ^16^ reported for Romney sheep that breeding for the short-tail phenotype possibly reduced the viability of embryos that were homozygous carriers of some putative short-tail alleles. In Merinos, James, et al. ^14^ observed increased incidences of rear-end defects. Zhi, et al. ^17^ discovered a c.G334T mutation in the *T* gene in the native Chinese Hulunbuir breed and showed that the T allele leads to the extreme short-tailed phenotype, i.e. tailless animals with exposed anus. To prove the causality of this mutation, they genotyped 120 short-tailed Hulunbuir sheep. The observed frequencies of the genotypes (17 G/G, 103 G/T and no T/T) are consistent with the embryonic lethality due to the T/T genotypes in *T* gene ^17^. A comparable association between short tails and embryonic lethality or malformations has been demonstrated in various breeds of dogs and cats too ^18–20^. These undesirable negative side effects discouraged and slowed down active breeding programs against overlong tails in the economically most important wool sheep breeds. Moreover, there have been no successful genetic mapping studies in long-tailed Merino sheep breeds, and the possible relationship between the short tail phenotype and embryonic viability or hind end malformations has never been investigated on a genetic basis in Merinos.

The aim of the present study was therefore:

1. To investigate the phenotypic and additive genetic variance in tail length in the Merinolandschaf, which belongs to the economically most important long-tailed Merino breed group worldwide;
2. To map the position of the major QTL(s) affecting tail length;
3. To detect and confirm causal candidate genes by sequencing and genotyping;
4. To determine the distribution of ancestral and derived alleles in a wide range of domestic sheep breeds with different tail lengths as well as in different wild sheep subspecies;
5. To contribute to the understanding of the relationship between genotype and phenotype during embryonic patterning and early development;
6. To put causal alleles in the evolutionary context of sheep species;

Together these objectives will enable more efficient breeding towards the ancestral phenotype and thus improve animal welfare in sheep production without negative side effects.

## Materials and Methods

### Animal samples and phenotypes

The entire mapping design of 362 phenotyped and genotyped animals were collected in three phases: (i) 236 Merinolandschaf lambs with very short (104) or long (132) tails were selected for phenotyping and sampling from 2293 visually inspected lambs on a farm in Lower Bavaria with no custom of tail docking, (ii) 102 random male lambs from the same farm were phenotyped and sampled, i.e. without preselection by visually inspected of tail-length and (iii) 24 Merinolandschaf lambs with short (19) or long (5) tails were selected from 102 visually inspected lambs by the Justus Liebig University of Giessen (JLU) (**Table 1**).

**Table 1:** Total number of visually inspected and sampled lambs at both sampling locations regarding to its phenotype

| Sampling location |  | Visually inspected | Sampled |  |  |  |
| --- | --- | --- | --- | --- | --- | --- |
|  |  |  | Total | Short | Long | Random |
| Lower Bavaria | 1. sampling | 2293 | 236 | 104 | 132 | - |
|  | 2. sampling | 110 | 102 | - | - | 102 |
| JLU Giessen |  | 102 | 24 | 19 | 5 | - |
| Total |  | 2505 | 362 | 123 | 137 | 102 |

Phenotyping of these 362 lambs was conducted according to the method proposed by Eck, et al. ^21^. Although an age of 5 weeks proved to be the optimal time point for phenotyping, we sampled and phenotyped also younger and older lambs in order not to disturb the work processes on the farm. Tail length (TL) was measured with a custom-made wooden board from the anus to the tip of the tail, body weight (BW) with a standard scale, and height at the withers (WH) with a metal measuring device from dog sports from the floor to the highest point of the withers. Furthermore, gender, age, and litter size were recorded. Unfortunately, the age and litter size are only approximately known for randomly sampled 102 animals. To improve haplotype inference, we sampled and genotyped 22 putative fathers of the above lambs. These rams were not phenotyped and contributed only indirectly to the QTL mapping.

All blood samples were taken according to best veterinary practice and under a permit from the Government of Upper Bavaria (permit number: 55.2-1-54-2532.0-47-2016), or the Regional Council of Gießen, Hassia (KTV number: 19 c 20 15 h 02 Gi 19/1 KTV 22/2020).

### Genotypes

All 362 phenotyped lambs and 22 sires were genotyped using Illumina’s OvineSNP50 BeadChip (Illumina, San Diego, USA) according to the manufacturer’s specifications. All physical marker positions were determined on the ovine reference genome assembly Oar_4.0 (https://www.ncbi.nlm.nih.gov/assembly/GCF_000298735.2) and the positions of all markers or sequences in materials, results or discussed below correspond to the Oar_4.0 reference genome unless otherwise stated. The chip contains 54,241 SNPs almost evenly distributed across the genome with an average marker spacing of 50.9 kb ^22^. Not all of these 54,241 SNPs were used for mapping. We filtered SNPs according to the following exclusion criteria: (i) unsuccessful genotyping in more than 5 % of the animals, (ii) frequent paternity conflicts in animals with known paternity, (iii) unknown position in the reference genome, (iv) minor allele frequency (MAF) of less than 0.025, and (v) localization on a sex chromosome since the analyses were exclusively carried out on autosomes. As a result, 45,114 markers remained in the marker set for the mapping analyses. Haplotype phasing and imputation were conducted using a Hidden Markov Model (HMM) implemented in *Beagle* version 5.0 ^23^. To improve the accuracy of haplotyping and imputation, genotype and pedigree information from about 5,100 additional animals genotyped with the OvineSNP50 BeadChip but not phenotyped were added to the haplotyping design ^24^.

### Estimation of heritability and mapping

First, we tested the heritability of the tail length in the pure Merinolandschaf breed. For this purpose we used the software *GCTA* v1.93.2, which has been extended with GRM, a tool for estimating the genetic relationship matrix and a genomic-relatedness-based restricted maximum-likelihood approach (GREML), to estimate the proportion of variance in our phenotype explained by all SNPs (the SNP-based heritability) ^25^. The sex, age, weight, and withers height of the lambs at phenotyping were modeled as fixed effects.

### Mixed linear model-based association analysis

To map a putative tail length locus, we performed mixed linear model-based association (MLMA) analyses with a leave-one-chromosome-out (LOCO) approach (Model 1) as implemented in the software *GCTA* v1.93.2 ^26^. Here we used the following model:

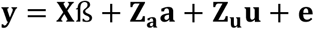

where **y** is the vector of tail lengths (cm), **ß** is a vector of fixed effects including the mean, sex, age, body weight (BW) and withers height (WH) at phenotyping, **a** is the vector of the additive effect (fixed) of the candidate SNP to be tested for association, **u** is the vector of polygenic effect (random or accumulated) of all markers, excluding those on the chromosome which contains the candidate SNP and **e** is a vector of residuals. **X**, **Za** and **Zu** are the incidence matrices for **ß**, **a** and **u**, respectively.

The suggestive significance threshold was set at *P* < 1/*N* and the genome-wide significance threshold at *P* < 0.05/*N*, according to Bonferroni method, *N* stands for the number of markers ^27^. For initial MLMA-analysis we considered 45,114 markers resulting in the suggestive *P*-value at 2.22 x 10^-5^ and genome-wide at 1.11 x 10^-6^.

A second MLMA (Model 2) analysis included one additional locus, which we detected during our research, as a consequence, we considered 45,115 markers. The suggestive and genome-wide significance threshold remained the same.

### Combined linkage disequilibrium and linkage analysis (*cLDLA*)

Parallel with SNP-based association analyses using MLMA, we performed multiple haplotype-based *cLDLA* analyses ^28^, which have been successfully used for binary and quantitative trait mapping in previous studies ^29–32^.

To correct for familial relationships and population stratification, unified additive relationships (UARs) were estimated between all animals on a genome-wide level ^33^. The inverse of the UAR matrix was then included in the variance component analysis. To account for linkage disequilibrium in the form of local haplotype relationships, the Locus IBD (LocIBD) was estimated according to the method of Meuwissen and Goddard ^34^ using sliding windows of 40 SNPs. For each window, we estimated LocIBD in the middle, i.e. between SNPs 20 and 21, based on the flanking marker haplotypes. Following the method for additive genetic relationship matrices (**G_RM_**) by Lee and Van der Werf ^35^, the matrix of LocIBD-values was converted into a diplotype relationship matrix (**DRM**).

Variance component analyses were carried out with the program *ASReml*^36^ according to the method of Meuwissen, et al. ^28^. *ASReml* estimated the maximum likelihood, variance components, and fixed and random effects simultaneously by considering the genome-wide UAR as well as the locus-specific (**DRM**) relationships matrices in the following mixed linear model:

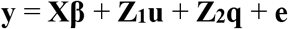

where **y** is again the vector of tail lengths and **β** is the vector of fixed effects (BW in kg, WH in cm, sex, age, and the overall mean μ; BW and WH data were both standardized and centered). The vector **u** is the vector of random polygenic effects (with 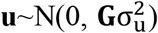) where **G** represents the matrix of genome-wide IBD estimations), **q** is the vector of random additive-genetic QTL effects (with 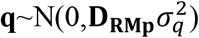) where **D_RM_p__** is the diplotype relationship matrix at position *p* of a supposed QTL), and **e** is the vector of random residual effects (with 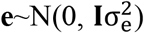), where **I** is an identity matrix). It is assumed that **u**, **q** and **e** are not correlated. **X, Z_1_, Z_2_** are incidence matrices linking the observed values with the fixed and random QTL effects.

The variance components and likelihood estimated with *ASReml* were then used in a likelihood ratio test statistic (*LRT*). The *LRT* values follow a *χ*^2^ distribution with one degree of freedom ^37^ and were calculated as:

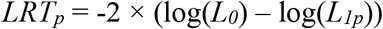

where log(*L_0_*) is the logarithm of the likelihood estimated by *ASReml* for the model without and log(*L_1p_*) with QTL effects at the center of the window *p*.

To obtain a significance threshold, a Bonferroni correction was carried out to account for multiple testing due to the 40-SNP sliding windows ^38^. This resulted in a corrected *P*-value of < 4.44 × 10^-5^ (i.e., 0.05/1127 where 1127 is the number of nonoverlapping 40 SNP windows) and a corresponding *LRT* value with genome-wide significance is equal to 16.67.

In addition to Model 3 described above, we performed two further genome-wide cLDLA analyses, with both analyses using candidate locus genotypes for the same set of animals. Model 4 contains only one additional locus and is therefore comparable to Model 2 of MLMA. In Model 5, the genotypes of the same candidate locus are considered as a fixed effect, i.e. **β** is the vector of μ, sex, age, BW, WH, and candidate locus effects. A comprehensive overview of the different models is provided in **Table 2**. For all maxima of the LRT curve (*LRT*max) that exceeded the genome-wide significance threshold, the 2-LOD (logarithm of the odds) criterion was used to determine the associated confidence intervals ^39^. Closely located LRT peaks were assumed to belong to the same QTL as described by Müller et al ^32^.

**Table 2:** Mixed linear models used for association and combined linkage disequilibrium and linkage analysis. The fix and random effect are the overall mean (μ), sex, age, body weight (BW), withers height (WH), the vector of the additive effect of the candidate marker to be tested for association (a), the vector of random polygenic effects (u), the vector of random additive-genetic QTL effects (q) and the vector of random residual effects (e)

| Analysis | Model name | Effects |  | Comment |
| --- | --- | --- | --- | --- |
|  |  | Fix | random |  |
| MLMA | Model 1 | $\mu$, sex, age, BW, WH, a | u, e | |
| | Model 2 | $\mu$, sex, age, BW, WH, a | u, e | add one candidate locus as marker |
| cLDLA | Model 3 | $\mu$, sex, age, BW, WH | u, q, e | |
| | Model 4 | $\mu$, sex, age, BW, WH | u, q, e | add one candidate locus as marker |
| | Model 5 | $\mu$, sex, age, BW, WH, cSNP, a | u, q, e | add one candidate locus as fix effect |

The results of the association analyses were presented as Manhatten plots produced by the *R* package *QQMAN* ^40^.

### Annotation of gene content and gene set enrichment analysis

The confidence intervals of the *LRT*max were compared with a map of annotated genes in the UCSC Genome Browser Oar_v4.0/oviAri4 Assembly ^41,42^ using the “RefSeq Genes” track. We refer to the Ensembl database of (Oar_v3.1) ^43^, the Mus musculus Assembly (GRCm39) ^44^, and the Homo sapiens Assembly (GRCh38.p13) ^45^ from NCBI ^46^ as well to consider genes encompassing the confidence intervals. Next, gene set enrichment analyses were carried out with the software *Enrichr* (Ontologies, MGI Mammalian Phenotype Level 4 2019) ^47,48^.

### DNA extraction, sequencing and analysis of the sequences

Genomic regions where the LRT curve reached a maximum value above the genome-wide significance threshold were sequenced using targeted capture sequencing on 48 selected lambs. To minimize the risk of missing some causal variations that are close but outside the confidence intervals, we increased the candidate regions on both sides by about 300 kb. This resulted in the captured regions on OAR2 from 93.200.000 to 96.700.000bp and on OAR11 from 36.600.000 to 37.900.000 bp. Genomic DNA was extracted from the blood samples using the ReliaPrep™ Blood gDNA Miniprep System.

Whole-Genome sequencing libraries were prepared from 250 ng of genomic DNA by tagmentation with the NexteraFlex kit from Illumina. Subsequently, the libraries were dual-barcoded and amplified by PCR, purified with SPRI beads and pooled in equimolar amounts. The pooled libraries were enriched for the target regions by hybridization to an Agilent capture array with 244k oligo spots. The oligo probes were selected from the repeat-masked DNA sequence as all possible 60mers that do not overlap with repeat-masked bases and that are staggered in 15 nt tiling steps concerning their neighbors. After 65 hours of hybridization in the presence of cot-I sheep DNA and adapter blocking oligos at 65°C the capture array was washed and the captured library molecules were eluted at 95°C for 10 min in a volume of 500 μl DNA-grade water. The enriched libraries were then amplified by PCR, analyzed on the Bioanalyzer and sequenced in 2*110 bp paired-end mode on a P2 flowcell of a NextSeq1000 sequencer from Illumina.

*Sickle* ^49^ was used to trim the adaptors and filter low-quality sequences of the raw reads in the FASTQ files. With *FastQC* ^50^, we assessed the quality parameters of the filtered sequencing data. The filtered reads were mapped to the sheep reference genome OAR_v4.0 using the default parameters of *BWA-MEM* ^51^ alignment tool. To convert the SAM files into coordinate sorted BAM files and to remove the duplicated reads, *Samtools* ^52^ and *Picard* ^53^ were used. Base quality recalibration and indel realignment were done with *GATK* ^54^. Variant calling was performed with *Bcftools* (mpileup) ^55^ for SNPs. To detect indels and structural variants (SV) we used *smoove* (*Lumpy*)^56^ and *delly*^57^ with default parameters.

To ensure that we do not miss any candidate variant we performed a visual examination of the captured regions using *jbrowse* ^58^, focusing on the region our candidate gene is located. We also examined the genetic variance in open source WGS of three sheep groups: 1) long-tailed domestic sheep, 2) short-tailed domestic sheep and 3) short-tailed Asiatic mouflon. **Table 3** shows the examined breeds/species including the Run-numbers and Biosample IDs.

**Table 3:** Run numbers and BioSample ID of different sheep breeds and there genotype for both candidate variants. Genotypes are homozygous ancestral (A/A), homozygous derived (D/D) and heterozygous (A/D)

| Breed | Sample |  | Genotype |  | Submitted by |
| --- | --- | --- | --- | --- | --- |
|  | Run number | BioSample ID | SV | SNP |  |
| Asiatic Mouflon | ERR157938 | SAMEA2012637 | A/A | A/A | Genoscope * |
| Asiatic Mouflon | ERR157930 | SAMEA2012638 | A/A | A/A | Genoscope * |
| Asiatic Mouflon | ERR157939 | SAMEA2012639 | A/A | A/D | Genoscope * |
| Asiatic Mouflon | ERR157942 | SAMEA2012640 | A/A | A/D | Genoscope * |
| Asiatic Mouflon | ERR157931 | SAMEA2012641 | A/A | D/D | Genoscope * |
| Asiatic Mouflon | ERR157932 | SAMEA2012642 | A/A | A/A | Genoscope * |
| Asiatic Mouflon | ERR157944 | SAMEA1967031 | A/A | A/A | Genoscope * |
| Asiatic Mouflon | ERR157935 | SAMEA2012643 | A/A | A/A | Genoscope * |
| Asiatic Mouflon | ERR332589 | SAMEA2065600 | A/A | A/A | Genoscope * |
| Asiatic Mouflon | ERR332575 | SAMEA2065601 | A/A | D/D | Genoscope * |
| Asiatic Mouflon | ERR332587 | SAMEA2065602 | A/A | A/A | Genoscope * |
| Asiatic Mouflon | ERR332582 | SAMEA2065603 | A/A | A/A | Genoscope * |
| Asiatic Mouflon | ERR332573 | SAMEA1972234 | A/A | A/D | Genoscope * |
| Asiatic Mouflon | ERR315509 | SAMEA2065604 | A/A | A/A | Genoscope * |
| Asiatic Mouflon | ERR466546 | SAMEA2395410 | A/A | A/A | Genoscope * |
| Asiatic Mouflon | ERR466544 | SAMEA2395411 | A/A | A/A | Genoscope * |
| Finnsheep | SRR11657543 | SAMN14590314 | A/A | A/A | Li, et al. <sup>89</sup> |
| Finnsheep | SRR11657544 | SAMN14590313 | A/D | A/D | Li, et al. <sup>89</sup> |
| Finnsheep | SRR11657545 | SAMN14590312 | A/A | A/A | Li, et al. <sup>89</sup> |
| Finnsheep | SRR11657546 | SAMN14590311 | A/A | A/A | Li, et al. <sup>89</sup> |
| Romanov | SRR12396891 | SAMN15517583 | A/A | A/D | Deng, et al. <sup>90</sup> |
| Romanov | SRR4291219 | SAMN05216760 | A/A | A/A | Heaton, et al. <sup>91</sup> |
| Romanov | SRR4291223 | SAMN05216759 | A/A | D/D | Heaton, et al. <sup>91</sup> |
| Romanov | SRR4291160 | SAMN05216766 | A/A | A/A | Heaton, et al. <sup>91</sup> |
| Swiss White Alpine | ERR3086436 | SAMEA5239874 | D/D | D/D | University of Bern** |
| Swiss White Alpine | ERR3086440 | SAMEA5239878 | D/D | D/D | University of Bern** |
| Swiss White Alpine | ERR3086476 | SAMEA5239914 | A/D | A/D | University of Bern** |
| Swiss White Alpine | ERR3086477 | SAMEA5239915 | A/D | A/D | University of Bern** |
| Rambouillet | SRR4291242 | SAMN05216757 | D/D | D/D | Heaton, et al. <sup>91</sup> |
| Rambouillet | SRR4291257 | SAMN05216755 | *** | D/D | Heaton, et al. <sup>91</sup> |
| Rambouillet | SRR4291268 | SAMN05216753 | D/D | D/D | Heaton, et al. <sup>91</sup> |
| Rambouillet | SRR6305143 | SAMEA104496890 | A/D | A/D | Baylor College of Med. |
| Ancient DNA Seq | ERR3861593 | SAMEA6516192 | *** | A/A | Yurtman, et al. <sup>63</sup> |
| Ancient DNA Seq | ERR3861592 | SAMEA6516191 | *** | A/A | Yurtman, et al. <sup>63</sup> |
| Ovis ammon | SRR8560952 | SAMN10915547 | A/A | A/A | CAAS**** |
| Ovis ammon | SRR8560953 | SAMN10915548 | A/A | A/A | CAAS**** |
| Ovis ammon | SRR9222805 | SAMN11979390 | A/A | A/A | CAAS**** |
| Ovis ammon | SRR9222806 | SAMN11979389 | A/A | A/A | CAAS**** |
| Ovis ammon | SRR9222807 | SAMN11979391 | A/A | A/A | CAAS**** |
| Ovis canadensis | SRR501858 | SAMN01000748 | A/A | A/A | Baylor College of Med. |
| Ovis canadensis | SRR501895 | SAMN01000746 | A/A | A/A | Baylor College of Med. |
| Ovis canadensis | SRR501898 | SAMN01000747 | A/A | A/A | Baylor College of Med. |
| Ovis dalli | SRR501847 | SAMN01000785 | A/A | A/A | Baylor College of Med. |
| Ovis dalli | SRR501897 | SAMN01000764 | A/A | A/A | Baylor College of Med. |
| Ovis vignei | ERR454945 | SAMEA2358291 | A/A | A/A | Genoscope * |
| Ovis vignei | ERR454946 | SAMEA2358287 | A/A | A/A | Genoscope * |
| Ovis vignei | ERR454947 | SAMEA2358290 | A/A | A/A | Genoscope * |
| Ovis vignei | ERR454948 | SAMEA2358289 | A/A | A/A | Genoscope * |
| Ovis vignei | ERR454950 | SAMEA2358291 | A/A | A/A | Genoscope * |
| Ovis nivicola | ERR4161992 | SAMEA6833340 | A/A | A/A | Upadhyay, et al. <sup>92</sup> |
| Ovis nivicola | ERR6667562 | SAMEA8657697 | A/A | A/A | Upadhyay, et al. <sup>93</sup> |
| Ovis nivicola | ERR6668200 | SAMEA8657699 | A/A | A/A | Upadhyay, et al. <sup>93</sup> |
| Ovis nivicola | ERR6668794 | SAMEA8657700 | A/A | A/A | Upadhyay, et al. <sup>93</sup> |
| Ovis nivicola | ERR6667561 | SAMEA8657698 | A/A | A/A | Upadhyay, et al. <sup>93</sup> |
| Ovis nivicola | ERR5858461 | SAMEA8657696 | A/A | A/A | Upadhyay, et al. <sup>93</sup> |

\* Sequenced as part of the NextGen project
\*\* Institute of Genetics
\*\*\* No reads mapped
\*\*\*\* Institute of Animal Science of CAAS

### Validation of candidate SNP using PCR

For one detected candidate SNP we performed genotyping by PCR-RFLP and electrophoresis on a 2% agarose gel on all 362 sampled lambs and there 17 confirmed fathers. The PCR primer sequences designed by *Primer3* ^59^ were TTTAAAACGCTTTGGATT (Forward, Left Primer) and CACTCGGCAGGAGTAGTA (Reverse, Right Primer). The used restriction enzyme was *BsrI.* It recognizes the mutant sequence TGAC/CN, where the G is the variable base and the “/” presents the site where cutting is performed. DNA amplification was performed with 35 cycles. The total reaction mixture was 15.0 μl containing 3.0 μl 5X buffer, 1.5 μl dNTPs (10mM), 0.6 μl of 10 μM Forward and Reverse Primer respectively, 1.0 μl DNA (15 ng/μl), 0.07 μl GoTaq®G2 DNA Polymerase (Promega, Madison, Wisconsin, USA) and distilled water.

We used 1.5 U of the enzyme *BsrI,* 3.0 μl DNA (PCR product), 2 μl Cut Smart Buffer, and distilled water for a total reaction volume of 20 μl. The reaction mixture was afterward incubated for 3 h at 65 °C. In the final step, we separated the DNA fragments by size and visualized them by GelRed™-stained agarose gel electrophoresis. Only sequences harboring the derived allele (SNP G) were cut, the two resulting fragments had a length of 120 bp and 259 bp. The sequence with the ancestral allele (SNP C) retained its length of 379 bp.

### Validation of candidate insertion using PCR and Sanger sequencing

We also performed genotyping by PCR and electrophoresis on a 2% agarose gel on the same 379 sampled sheep for one detected causal candidate insertion. Multiple PCR primer sequences designed by *Primer3* ^59^ were tested (Supplementary Table S1), those that worked best are TTTATGAGCTTCTCTCCGCCA (Forward, Left Primer) and <u>CACTCGGCAGGAGTAGTA</u> (Reverse, Right Primer). DNA amplification was performed with 35 cycles. The total reaction mixture was 25.0 μl containing 5.0 μl 5X buffer, 2.5 μl dNTPs (10mM), 1 μl of 10 μM Forward and Reverse Primer respectively, 1.0 μl DNA (15 ng/μl), 0.07 μl GoTaq®G2 DNA Polymerase (Promega, Madison, Wisconsin, USA) and distilled water. In the final step, we separated the amplicons by size and visualized them by GelRed™-stained agarose gel electrophoresis.

Two lambs, which are homozygous for the SV, one lamb, which is homozygous for the ancestral allele and two heterozygous lambs were resequenced using Sanger sequencing with the above mentioned best working Primers. The amplicons were sequenced using the cycle sequencing technology (dideoxy chain termination / cycle sequencing) on ABI 3730XL sequencing machines (Eurofins Genomics, Germany). The sequenced data were analyzed using *SnapGene* software (from Insightful Science; available at https://www.snapgene.com/).

## Results

### Initial mixed linear model association analysis

The *GCTA-GREML* analysis revealed SNP-based heritability of 0.992 (standard error of 0.12), meaning that a very high proportion of the tail length variance in Merinolandschaf breed is explained by genome-wide SNP markers. Despite very high heritability, i.e., close to 1, the initial association analysis (MLMA Model 1) revealed no genome-wide significant association between any SNPs and tail length. Even the four most significant SNPs (**Figure *1*a**) remain below the suggestive significance threshold of *P* = 2.22 x 10^-5^.

**Figure 1:**
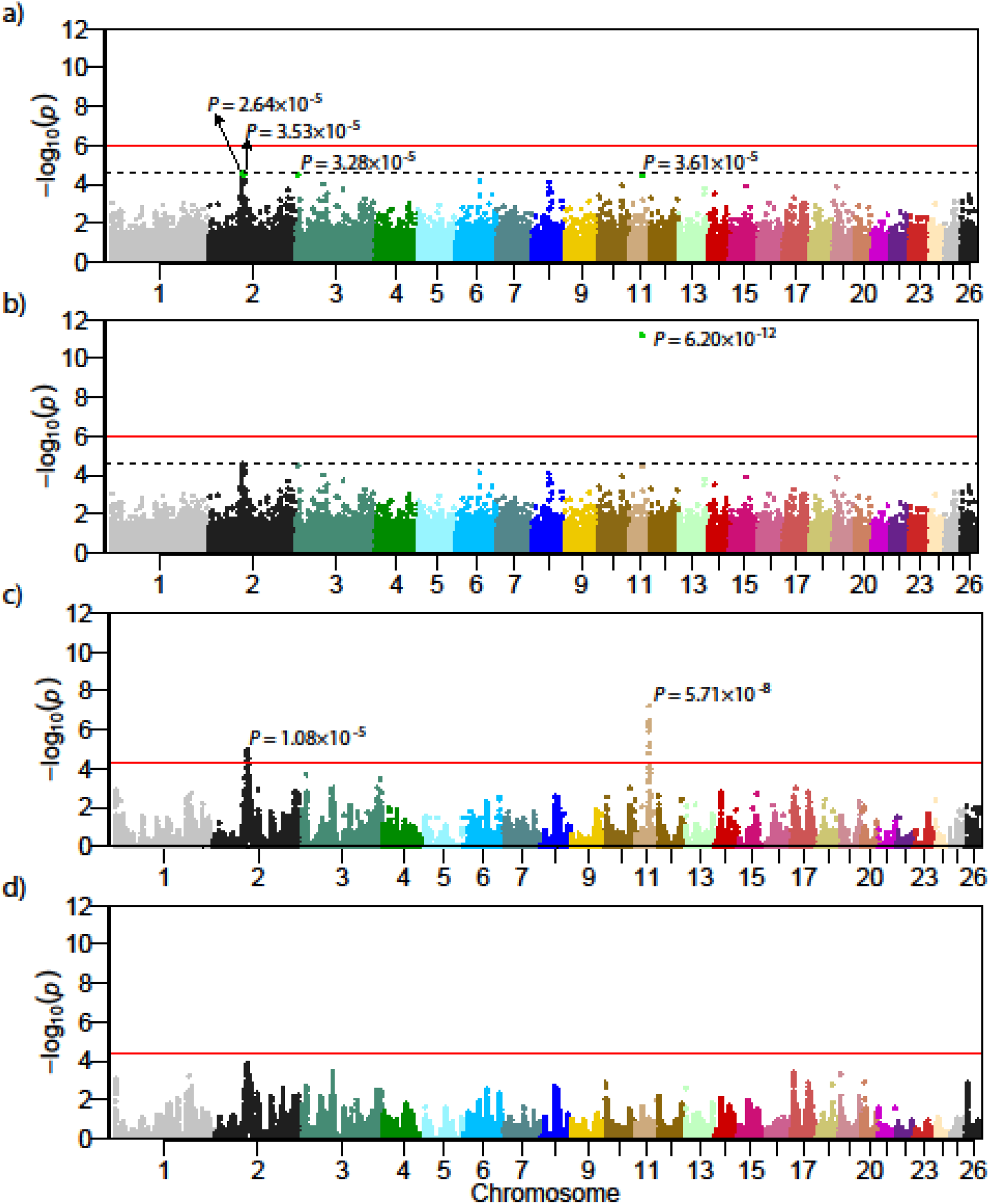
Results of performed mixed linear analyses presented as Manhattan Plots; (a) MLMA Model 1 with 45,114 markers no –log10(*p*-value) were above the suggestive line, the four markers with the lowest *p* values are shown; (b) MLMA Model 2 with 45,115 markers, the additionally added candidate locus on OAR11 shows genome-wide significance; (c) cLDLA Model 3 with 45,114 markers resulted in two genome-wide significant peaks on OAR2 and OAR11; (d) cLDLA Model 5 with 45,115 markers and candidate locus added as fix effect, the peak on OAR2 decreases below the genome-wide significance and the peak on OAR11 erases completely.

### Initial combined linkage disequilibrium and linkage analysis

The haplotype-based cLDLA mapping (cLDLA Model 3) resulted in two genome-wide significant QTLs associated with tail length in Merinolandschaf (**Figure *1*c**). The most prominent and narrow peak is on the sheep chromosome 11 (OAR11) at position 37,111,462 bp with *LRTmax* = 29.460 corresponding to *P* = 5.71*10^-8^ (Bonferoni corrected: *P* = 6.43*10^-5^). The second genome-wide significant QTL affecting tail length was mapped to the chromosome 2 at position 94,538,115 bp with *LRTmax* = 19.356 corresponding to *P* = 1.08*10^-5^ (Bonferoni corrected: *P* = 1.22*10^-2^).

Applying the 2-LOD criterion, the corresponding confidence interval was set for the *LRTmax* on OAR11 between positions 37,000,925 bp and 37,521,490 bp and for the maximum value on OAR2 between positions 93,441,900 bp and 96,402,884 bp. These intervals were then considered in the UCSC Genome Browser Oar_v4.0/oviAri4 Assembly, which resulted in the list of genes summarized in **Table *4*** and **Table *5***. The list of positional candidates also includes obvious functional candidates from the sheep homeobox B gene cluster (Chr11:37,290,203-37,460,240) with *HOXB13* (37,290,203-37,292,513) as the closest and most prominent candidate, lying only 179-Kb proximal to the *LRTmax*.

**Table 4:**
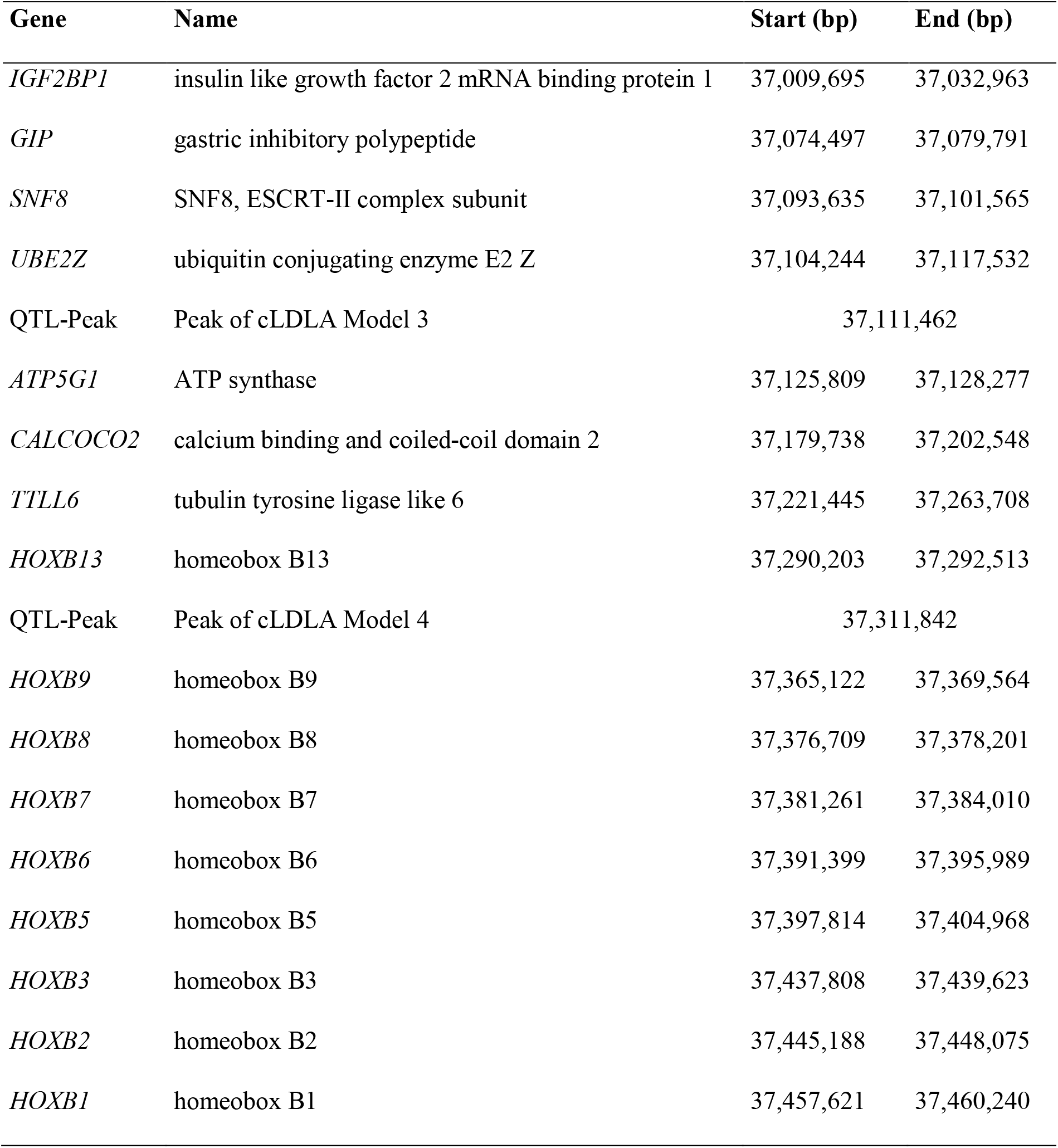
Genes on OAR11 between positions 37,000,925 bp and 37,521,490 bp as well as the genome-wide significant peaks of cLDLA Model 3 and 4

**Table 5:**
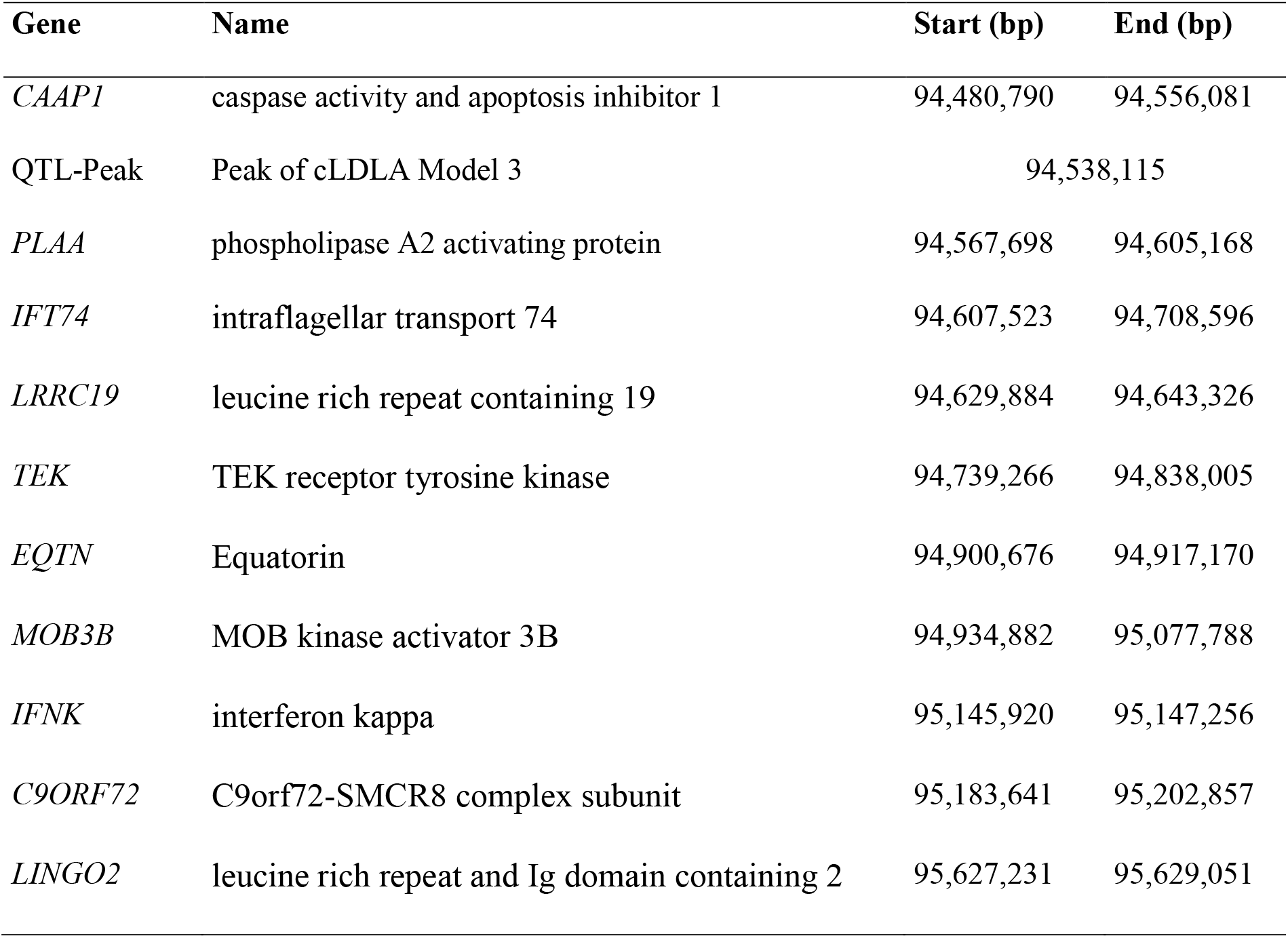
Genes on OAR2 between positions 93,441,900 bp and 96,402,884 bp as well as the genome-wide significant peak of cLDLA Model 3

Additional chromosome-wide significant peaks were observed on OAR2, OAR3, OAR10, OAR14, and OAR17. However, these peaks show *LRT* values far below the genome-wide significance and thus, were not investigated further.

### Estimation of QTL effects and selection of animals for capture sequencing

In the previous step of cLDLA, we used *ASReml* to estimate variance components, fixed and random effects affecting tail length in Merinolandschaf breed. Here, we analyzed in more detail the estimated effects at loci with the most significant association, i.e. at loci showing *LRTmax* values. We sorted all 362 lambs according to the random additive genetic QTL effects (vector **q**) estimated at *LRTmax.* Together with the QTL effects, we simultaneously considered all input (**y**, sex, age, WH, BW, maternal and paternal haplotypes at 40-SNP window with *LRTmax* in interval between SNP 20 and 21) and output data (**u**, **ß** and **e**) that contributed to *LRTmax.* Visual inspection of this table allowed us to select 48 lambs (half with the most negative and half with the most positive additive genetic QTL effects) for targeted capture sequencing. Supplementary Figure S1 and S2 show the distribution of the sequenced lambs regarding tail length and diplotype effects on OAR2 and OAR11

A regression analysis performed with the function *lm* in *R*^60^ estimated the adjusted coefficient of determination of *R^2^*=0.58 for the *LRTmax* on OAR11 and only *R^2^*=0.15 for the *LRTmax* on OAR02 when using tail length as the dependent and diplotype effect as the independent variable. Adding, age, sex, BW and WH as additional independent variables yield *R^2^*=0.78 for QTL on OAR11 and *R^2^*=0.45 for QTL on OAR02 (Supplementary Table S2 and S3). According to the shape of the LRT curve, the significance of the mapping and the coefficient of determination, the haplotypes associated with the putative causative alleles are more distinct in QTL on OAR11 than on OAR02. However, the selection of 48 lambs for capture sequencing represents a trade-off between the two QTLs, with the choice for OAR11 being more decisive. The selected lambs could be divided into two groups: 23 long-tailed with positive QTL effect and 25 short-tailed lambs with negative QTL effect on OAR11. Sorting the same lambs by QTL effects on OAR2 changes the order within the group and results in 3 individuals from the long-tailed group and 4 individuals from the short-tailed group moving to the other group.

### Capture sequencing of 48 lambs and detection of candidate mutations

Capture Sequencing was carried out at a mean depth between 0.41 and 2.45 on the target region of OAR2 and between 1.01 and 2.39 on the target region of OAR11. This coverage is much lower than intended and most possibly caused by competition with WGS performed on the same sequencing lane. However, applying default settings of bcftools mpileup on the individual samples, we detected 27,256 and 74,485 SNPs respectively on OAR11 and OAR02. Next, we sought to identify variants showing significant differences in frequency between short-tailed and long-tailed groups. Interestingly, of all the detected genomic variations, only one SNP satisfied the frequency-based criterion; this SNP (C→G) was located on OAR11 at the position of 37,290,361. The visual examination using *Jbrowse* confirmed the base substitution (*rs413316737*) as a nonsynonymous point mutation within the first exon of *HOXB13* (relative position 23). The point mutation results in a p.(Thr8Ser) substitution. All sequenced Merino lambs from the long-tailed group are homozygous for this missense variant (G/G). In the short-tailed group, 4 lambs are homozygous G/G, 6 are heterozygous C/G and 15 are homozygous C/C on that position. The Ensembl Variant Effect Predictor ^61^ predicted a SIFT score of 0.54 and classified the mutation as so-called *“Tolerated”* missense variant.

In the next step, protein BLAST ^62^ was used to align the amino acid sequence of the mutant HOXB13 protein against the amino acid sequence of wild-type HOXB13 protein of different mammals including all the extant wild sheep species (**Table 3**). This cross-species alignment (**Figure *2***) revealed that the amino acid at which the variant discovered here occurred is conserved. In addition, 5 downstream amino acids are also conserved among here aligned species.

**Figure 2:**
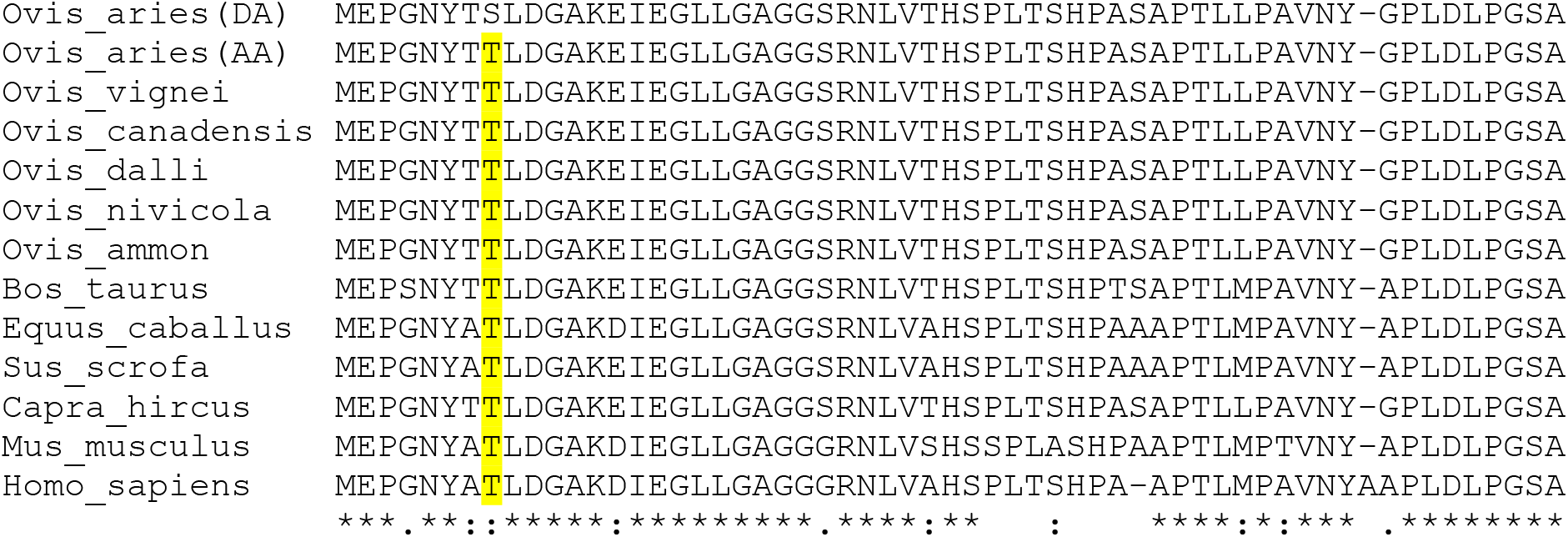
Amino acid sequence alignment of HOXB13 in multiple mammals. Positions of interest are highlighted. The asterisks present unique, the colons high similar and the single point moderate similar amino acids in every species on the respective position. To highlight the area of interest we used Clustal Omega ^87,88^ DA: Derived Allele, AA: Ancestral Allele

By comparing the allele frequency in long-tailed and short-tailed sheep in multiple breeds, we observed that the above-mentioned derived allele G occurs more frequently in long-tailed sheep breeds (Supplementary Table S4). Further, at this position, we only observed the ancestral allele in Urial, Argali, Snow sheep, Dall sheep, Canadensis and two ancient (~8,000 years) sheep genomes ^63^. On the other side out of 16 investigated Asiatic mouflon 3 were heterozygous C/G and 2 homozygous G/G for the point mutation. However, it is worth mentioning here that one Asiatic mouflon (G/G) and the two ancient WGS have low coverage (**Table 3**) at this locus.

To investigate the pattern of structural variations (SVs), we increased the sequencing coverage of both the groups by pooling the raw reads of their respective samples. Further, we identified SVs in the targeted region (between 36,600,000 and 37,900,000 bp) of OAR11 from these data by using the three different approaches: *Smoove, Delly* and visual examination using *Jbrowse*. Using the strict threshold criteria, i.e. SV not present in a pooled sample of short tail sheep but present in a pooled sample of long tail sheep, we identified 27 and 32 SVs from *Smoove* and *Delly* approaches, respectively. However, it is noteworthy that our captured sequencing had highly non-uniform and relatively low coverage, therefore, these approaches might have missed many true positive and included high number of false positives. In fact, our visual examination of these regions also indicated so (Supplementary Figure S3).

Interestingly, window-by-window visual examination of the targeted region in *Jbrowse* revealed two distinct clusters of reads showing soft-clipped **(Figure 3a**) just about ~130 bp upstream of the candidate SNP. Both these clusters were arranged next to each other; while one cluster was present in all the sequenced animals, indicating either assembly error or assembly-specific variant as the likely cause. Another cluster of soft-clipped reads had distinct frequency distribution between the groups of short-tailed and long-tailed reads. At position 37,290,229 on OAR11, the long-tailed group showed 34 of the 35 reads mapped as soft-clipped, while the short-tailed group showed 13 of the 54 reads as soft-clipped.

**Figure 3:**
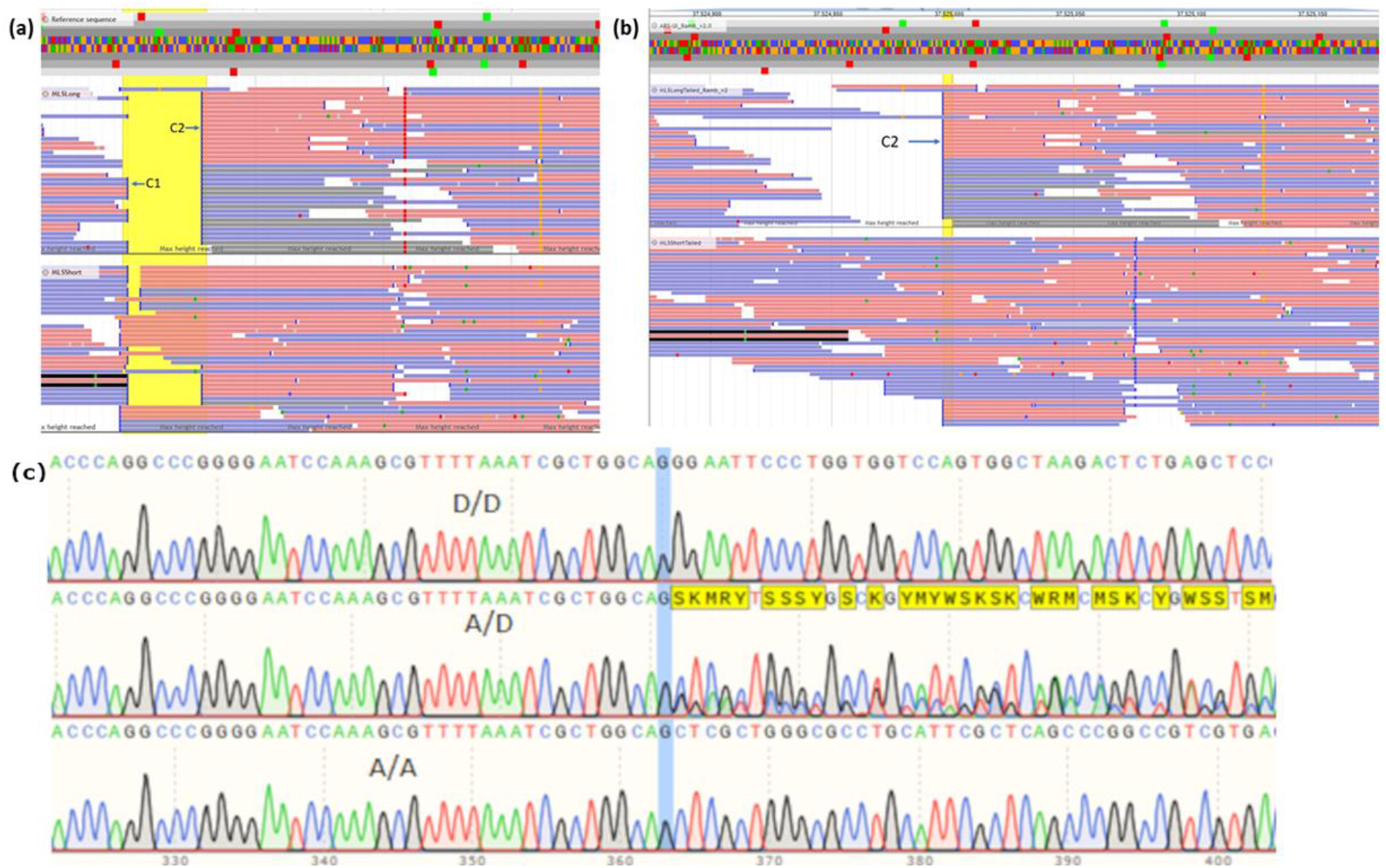
Screenshots showing insertions, position of the candidate SNP on 37,290,631 and extracts from the .ab1 trace files from sanger sequencing; a) Reads of the pooled sequenced long-tailed (above) and short-tailed (below) Merinolandschaf lambs mapped on assembly Oar_4.0, note the two clusters, cluster one is likely due to an assembly problem, cluster two represents an insertion shown in 34 of the 35 reads in the long-tailed group and in 13 of the 54 reads in the short-tailed group; b) shows the same groups mapped against the newest assembly ARS-UI_Ramb_v2.0, note the disappearance of cluster 1; c) sanger sequences for one homozygous derived (D/D), one heterozygous (A/D) and one homozygous ancestral (A/A) lamb around 40 bp before and after the breakpoint of the real insertion

We also observed a significant difference in the frequency of the clipped reads around this position between the WGS data of short and long-tailed sheep that were downloaded from NCBI SRA (**Table 3**). Therefore, in the next step, we aligned the pooled reads of the short-tailed and the long-tailed group and the WGS data of NCBI SRA against the latest sheep assembly (ARS-UI_Ramb_v2.0). Visual examination of the region upstream of the first exon of *HOXB13* gene on OAR11 revealed only one cluster of soft-clipped reads at positions 37,524,996 **(Figure 3b**) indicating that another cluster which was identified in the mapping against Oar 4.0 assembly was due to missing sequences of about 40 bp in Oar 4.0 assembly. Interestingly, we observed that a very high number of soft-clipped reads in this cluster had supplementary alignment on OAR5. Further, on OAR11 at the breakpoint, we also observed discordant alignment (overlapped) between forward and reverse reads. Based on the presence of the soft-clipped reads and discordant alignment, we suspected the presence of insertion or translocation. While we were investigating this region on ARS-UI_Ramb_v2.0 assembly, we came across a pre-print by Li, et al. ^64^; they reported insertion in the same region using the graph-assembly based method on the Pac-bio sequencing data of 13 different sheep breeds.

To investigate the soft-clipped regions further, we carried out Sanger sequencing of 5 samples, based on the previously described candidate SNPs. The analysis of the Sanger sequencing data **(Figure 3c**) and subsequent alignment against OAR11 in ARS-UI_Ramb_v2.0 using *BLAST* identified the SV as an insertion of 167 bp. Interestingly, we further observed that this sequence is flanked by 14 bp of direct repeats (CTGCCAGCGATTTA) on both sides. Therefore, we hypothesized that this insertion could be a part of SINE repeat family. Next, we searched for this sequence in DFAM repeat database and identified it as belonging to OviAri-1.113 SINE family.

### Genotyping of the most plausible candidates in 362 lambs and remapping of tail length

To confirm the association between the detected variants and tail length, we performed genotyping of these two candidates in the entire mapping population and used genotypes in the GWAS and cLDLA Model 2, 4 and 5 (**Table 2**). The PCR-genotyping of the candidate SNP resulted in 220 G/G, 118 C/G and 24 C/C lambs. The PCR-genotyping of the 132 bp upstream candidate insertion of 167 bp showed an identical distribution of genotypes over entire mapping population, i.e. insertion occurred in all haplotypes harboring the base G and never in haplotypes harboring C on the position 37.209.361. Due to the complete linkage between the insertion and the missense SNP, both candidates are considered synchronously and equally in Model 2 (MLMA) and Models 4 and 5 (cLDLA). The distribution of alleles in wild sheep subspecies and domestic sheep allow us to infer ancestral and derived alleles at both candidate loci. The absence of insertion and base C at c.C23G SNP of *HOXB13* are ancestral while 167bp insertion and base G are referred to derived alleles.

All lambs homozygous for ancestral alleles could be classified as short-tailed with mean tail-length 24.1 cm (±1.34) and mean QTL effect −2.92 cm (±0.71). On the other side lambs homozygous for derived alleles classified as long-tailed as well as short-tailed. Consequently, lambs with derived homozygous genotype show higher average tail length 31.5 cm and 4.15 times higher standard deviation of tail length (±5.56). Lambs with heterozygous genotype show average tail length of 25.7 and 2.73 times higher standard deviation (±3.66). The **Table 6** summarizes phenotype, QTL and polygenic effects of candidate insertion and SNP that are in population-wide linkage disequilibrium.

**Table 6:** The number of individuals, mean and SD of tail length, QTL effects and polygenic effects of the different genotype groups. The groups are homozygous ancestral (A/A), homozygous derived (D/D) and heterozygous (A/D)

| Genotype | Count |  |  | Tail length |  | QTL effect |  | Polygenic effect |  |
| --- | --- | --- | --- | --- | --- | --- | --- | --- | --- |
|  | all | short | Long | Mean | SD | Mean | SD | Mean | SD |
| D/D | 210 | 70 | 150 | 31.5 | 5.56 | 0.68 | 1.27 | 1.12 | 3.56 |
| A/D | 118 | 94 | 24 | 25.7 | 3.66 | -1.3 | 0.89 | -1.61 | 2.96 |
| A/A | 24 | 24 | 0 | 24.1 | 1.34 | -2.92 | 0.71 | -2.19 | 1.52 |

### *MLMA* and *cLDLA* including candidate variants as markers

To investigate the impact of derived alleles on the results of the SNP-based association analysis and the haplotype-based mapping we considered candidate locus as an additional marker located on Chr11:37,290,361 (see Model 2 and 4 **Table 2**). Thereby, only the number of considered markers increases by one compared to the original analysis, while the other input data, as well as the parameters and the model, do not change. This minimal change in the input data led to enormous changes in the results of the association analysis and limited changes in the results of the haplotype-based mapping. *MLMA* Model 2 confirmed the candidate causal locus as a uniquely genome-wide significant (*P* = 6.2 *10^-12^) locus, while *cLDLA* revealed a slightly altered significance (*LRTmax=29.112)* at the slightly altered position 37,311,842 bp. However, in the initial mapping *LRTmax* was 179 Kb away from *HOXB13,* and the Model 4 of *cLDLA* placed *LRTmax* between *HOXB13* and *HOXB9* (**Fehler! Verweisquelle konnte nicht gefunden werden.**). Therefore, the distance between *LRTmax* and the candidate gene *HOXB13* decreased from 179 Kb to 19 Kb, and thus *HOXB13* became the closest gene to *LRTmax.*

### Variance components, *MLMA* and *cLDLA* including candidates as fixed effect

The above results point to *rs413316737* and/or the insertion as plausible candidates for causal variations. The mixed linear model (MLMA or cLDLA) allows us to model important causal candidates and thus improve the mapping of residual variance (if present) in the mapping population. To investigate the presence of additional loci affecting tail length in long-tailed Merino sheep, we modelled the genotypes at the candidate insertion and SNP as fixed effects, i.e. lambs with homozygous ancestral genotype were classified as class 1, heterozygous as class 2 and homozygous derived as class 3, while the other input data, parameters and model did not change. We first estimated the SNP-based heritability of tail length after correcting for the most significant causal candidates. The heritability decreased from *h^2^*=0.992 to 0.898. This indicates still high proportion of the additive genetic variance in phenotyping variance corrected for the most plausible candidate mutations in *HOXB13.* However, as shown in **Figure 1d**, the cLDLA is unable to highlight additional candidates, although a relatively high proportion of additive genetic variance is still present in the mapping population studied here (*h^2^*=0.898). As expected from the true candidate, the mapping signal on OAR11 disappeared completely (**Figure 1d Figure *2***). Moreover, the LRT value at the peak on OAR02 decreased from 19.356 to 14.476 and marginally changed his position from 94.538.115 bp to 94.345.619 bp. With this change, a possible QTL on OAR02 loses its significance.

## Discussion

The present study aimed to identify the genes or variants responsible for the natural variability of tail length in the Merinolandschaf breed. The results presented here suggest that the inheritance of tail length depends largely on additive genetic variance almost without environment effects (*h^2^* =0.992). Despite the relatively small number (362) of phenotyped and genotyped animals, heritability was estimated with a relatively low standard error (SE=0.12). According to Visscher et al ^65^, the SE depended only on the sample size and became below 0.1 by using more than ~3000 independent individuals.

The GWAS (MLMA), i.e. the standard method for mapping loci associated with complex traits and diseases, showed no significant and even no suggestive association in our design with 362 animals and 45,114 markers. Typically, GWAS ^65^ uses several hundred thousand markers in large mapping designs. Therefore, the solution would be to enlarge the sample size and genotype this enlarged mapping design with high marker density (e.g. OvineHD). However, in studies such as this, where phenotypes are not routinely collected, the number of phenotyped animals may be limited or can only be expanded at great expense. On the other hand, the number of markers can also be a limiting factor in many species, e.g. there is only 50K chip for domestic goats. The alternative solution could be to apply a mapping method that uses more information from the current design. Due to time and cost constraints, we opted for haplotype-based mapping and obtained a highly significant (*P* = 5.71*10^-8^) and fine mapped QTL on OAR11 (CI 37,000,925 - 37,521,490 Mb) as well as another genome-wide significant result (*P* = 1.08*10^-5^ with less sharp mapping (OAR02, CI 93,441,900 - 96,402,884 Mb).

Previous research on tail length in domesticated animals, including sheep, mainly pointed to the *T* gene, also known as the brachyuria gene, as a candidate causal gene. In Hulunbuir short-tailed sheep, a *c.G334T* mutation in *T* gene is the main cause of the extreme short-tail phenotype ^17^ Mice that are heterozygous for mutations in the *T* gene have a short tail and homozygous embryos die in the middle of the gestation ^66–68^. In Manx cats, the short-tailed phenotype is caused by naturally occurring mutations in *T* gene ^19^, in particular by three 1-bp deletions. The *T* gene has also been associated with the short-tailed phenotype in various dog breeds ^18,69,70^. Additionally, *ANKRD11, ACVR2B* and *SFRP2* were detected as plausible candidate genes that could contribute to the reduction in tail length in particular dog breeds ^69^ and the somite segmentation-related gene *HES7* ^71^ in Asian domestic cats.

In this study we could not detect any increase in the LRT curve in the *T* gene region (OAR8: 87,717,306–87,727,483 bp). Furthermore, neither *ANKRD11* (OAR14: 13,810,611–13,882,911 bp), *ACVR2B* (OAR19: 11,794,562–11,802,479 bp) or *SFRP2* (OAR17: 3,727,698–3,736,241 bp) showed a significant increase in the LRT value. *HES7* is located on OAR11 (27,284,897-27,287,414 bp) but is 10 Mb away from the QTL confidence interval and was therefore not considered as a candidate gene for tail length in the Merinolandschaf breed. The confidence interval on OAR11 includes complete ovine *HOXB* cluster and additional seven genes (**Table 4**). Among these genes, a potential influence on tail length could be predicted for *HOXB6, HOXB8* and *HOXB13.* Here, *HOXB13* is the gene closest to the maximum *LRT* value and a corresponding literature search (see Diaz-Cuadros, et al. ^72^ for a review) indicates this gene as the most likely gene causing the large effect in our design. The literature search for plausible candidates for genes at OAR02 (**Table 5**) yielded hardly any useful results. We tried to improve our search with MGI Mammalian Phenotype level 4 (MMP4) ontology, a method for classifying and organizing phenotypic information related to mammalian species (66). After correction for multiple testing (adjusted *P* < 0.05), 29 ontologies were significantly enriched, but we see no plausible link to tail length.

To select the most suitable lambs for capture sequencing of confidence intervals of QTLs on OAR11 and OAR02 we mutually considered haplotypes, phenotypes, fixed effects and random effects at *LRTmax* of both regions. Again, the visual inspections, as well as linear regression analyses, confirmed QTL on OAR02 as inconclusive. This is evident from adjusted *R^2^* = 0.58 and only 0.15 for tail length fitted to QTL of OAR11 and OAR02, respectively.

Consistent with the clues in favor of OAR11 discussed above, capture sequencing revealed one plausible point mutation and one SINE insertion in the QTL region. All Merinolandschaf lambs and their fathers show a complete linkage disequilibrium between these two variants. This is not surprising because these mutations are only 132 base pairs away from each other, and recombination in such a short segment should be extremely rare. We further investigated this linkage in some typical long-tailed and short-tailed breeds (**Table 3**). The linkage was confirmed for long-tailed White Swiss Alpine and Rambouillet breeds but we observed the occurrence of the missense mutation without the insertion in some individuals of short-tailed domestic sheep breeds as well as in five Asiatic Mouflons (**Table 3**). Our analyses of the WGS data (**Table 3**) and amino acid sequence alignment (**Figure 2**) identifies allele C to be ancestral. Therefore, allele G is a derived but relatively ancient allele that segregates in *Ovis gmelini* and *Ovis aries*. The 167 bp insertion in the promotor region is also derived but more recent and segregates exclusively in domestic sheep and predominantly in long-tailed sheep breeds. This insertion occurred in the haplotype with the older missense allele G and both derived alleles segregate as a block. So, we did not observe the insertion without allele G, but we did observe the older allele G without insertion. The presence of the insertion in long-tailed sheep breeds and its location in the promotor region of *HOXB13* increases the probability of insertion as a causative variant for the long tail phenotype. Most likely because of its location the insertion modulates the promoter activity of *HOXB13* and leads to a longer tail by reducing the expression of *HOXB13* gene. Indeed, a recent study by Li, et al. ^64^ also detected an insertion of 169 bp close to the 5’ UTR of *HOXB13* at position 37,525,005 (ARS-UI_Ramb_v2.0) using the graph-assembly based method on the Pac-bio sequencing data of 13 different sheep breeds including Merino. Therefore, it is likely this study and Li, et al. ^64^ both identified the same insertion. The small differences in length and position of this insertion can be attributed to sequencing error due to the presence of long homopolymer of “T” base in the identified SINE repeat element.

Interestingly, the missense mutation (*rs413316737)* in the first exon of *HOXB13* (37,290,361 C→G) is included on the OvineHD array as SNP marker *oar3_OAR11_37337253.* This offers the possibility to check the allele distribution in available open source genotypes (Supplementary Table S4). It is noteworthy, that top-allele G of *oar3_OAR11_37337253* correspond to ancestral C in the coding sequence of the *HOXB13.* Genotyping of the candidate SNP and insertion throughout the mapping design provides the opportunity to test the efficiency of GWAS and cLDLA with the candidates as a marker and as a fixed effect. The inclusion of the candidates as a marker supports very strong association (i.e. *P* = 6.2 *10^-12^) or identity of the *rs413316737* and/or insertion with the causative locus. Additionally, it demonstrates the power of GWAS when the design includes causal variants or markers with population-wide LD. On the other hand, the inclusion of candidates as markers in haplotype-based mapping leads only to partial change, i.e. the mapping significance remains about the same, but the mapping peak is 9-times closer to the most plausible candidate gene. Even more conclusive is the impact of these mutations as a fixed effect in the model, because the correction for the true causal variant should cancel out the LRT peak on OAR11, and if this explains a full additive genetic variance, the heritability should also be reduced towards zero. Indeed, this model erase the LRT peak on OAR11 **(Figure 1d**) but the heritability remains relatively high (0.898). We thus gathered further evidence pointing to a missense mutation in the first exon of *HOXB13* and/or a structural variation in the promotor region of *HOXB13* as plausible causal mutations but also confirmed the presence of other, yet unknown causal variants that explain a large part of the phenotypic variance.

The *HOXB13* belongs to the family of homeobox genes, which were first described in *Drosophila melanogaster* ^73^. This gene codes for transcription factors and play an important role in structuring the body plan during embryogenesis (reviewed by Diaz-Cuadros, et al. ^72^). In mammals, there are 39 *Hox* genes organized in four clusters and 13 paralogous groups ^74^. There is functional redundancy among the paralogous *Hox* genes ^75^ and the paralog alleles can compensate for each other to a certain degree ^76^. Therefore, the loss of function of one *Hox* gene from a cluster is usually compensated by the functionality of an intact paralogue and only the loss of function of several paralogs results in more severe consequences in axial structuring ^74^. There is indirect evidence that *HOXB6* ^77^ and *HOXB8* ^78^ may influence tail length and embryonic viability. However, capture sequencing finds no genetic variants associated with tail length in any of these two candidate genes. Therefore, we do not discuss these positional candidates further.

As vertebrate embryos develop progressively from head to tail, the HOX13 paralogous group has been proposed to control axis termination ^72^, which is mainly achieved through regulation of proliferation and apoptosis activity in the posterior embryonic regions ^79^. In mice, loss-of-function mutations in *Hoxb13* lead to overgrowth of the spinal cord and caudal vertebrae in homozygous mice. These animals consequently show longer and thicker tails while viability and fertility remain unaffected ^79^. Premature expression of genes of the *Hox13* paralogous group, on the other hand, negatively influences the extension of the caudal axis and results in a truncated phenotype ^78^. Among all candidate genes from the *Hox* family, *HOXB13* is thus our best functional candidate gene. Moreover, Li, et al. ^64^ used previously published RNA-Seq data of sheep colon, performed a luciferase reporter assays and indicated an association of lower expression of *HOXB13* with insertion in its promotor region. This is in the line with above findings in mice. However, due to complete linkage, we cannot exclude the cooperative causality of the insertion and missense mutation in the exon 1. The *HOXB13* is expressed in the prostate of adult humans and is intensively studied as a candidate biomarker for the prognosis of prostate cancer (see Ouhtit, et al. ^80^ for review). These studies show that missense mutations in the coding sequence of *HOXB13* can change the affinity ^81^ or half-life ^82^ of heterodimer between HOXB13 and e.g. MEIS1 proteins. However, it was not within the scope of this study to investigate the affinity between different transcription factors in growing sheep embryos or cell lines. It is well known that the SINE insertion alone can alter gene expression in multiple ways ^83^, but we propose to test the hypothesis of a possible causality of the combination of altered expression and altered amino acid sequence.

As already introduced, attempts were made in 1970s and 1990s to breed short-tailed Romney ^16^ and Merino ^11,14^ sheep. These attempts failed due to evidence of reduced viability of presumably homozygous short-tailed Romney embryos and by increased incidences of rear-end defects in short-tailed Merinos. However, these observations contradict the fact that the short tail is the ancestral trait and that old Nordic short-tailed breeds like Romanov and Finnsheep are very viable and highly fertile. Therefore, we assume that these early experiments were not carried out with animals that had shorter tails due to ancestral genetic variants, but due to some recent deleterious variants. James et al ^14^ wrote that the tail lengths of the four sires measured before mating were all less than 5 cm. This is very short for an adult ram, even much shorter than the tail of short-tailed adult Romanov rams (~20 cm). Also, Romney sheep used for experimental mating by Carter ^16^ were described as “tailless”. Therefore, short or tailless phenotypes described in initial sheep breeding attempts ^14,16^ are comparable with deleterious mutations described for certain dog, and cat breeds ^18–20,69,70^ rather than to some ancestral alleles. In this study detected causal candidates for short tails are highly frequent or fixed in some very viable and highly fertile breeds (e.g. Finnsheep, Romanov, Dalapaels, Soay, Supplementary TableS2). Therefore, the selection toward ancestral allele do not carry any detrimental side effects for fertility or malformations ^84^ and can be achieved without introgression.

In our design, all homozygous ancestral and the majority (79.7%) of heterozygous lambs could be classified as short-tailed, while homozygous derived lambs are mostly (71.4%) but not always long-tailed (**Table 6**). This could be caused by interaction (epistasis) with alleles at other loci in the sheep genome or simply by the polygenic nature of the high proportion of additive genetic variance which was still not explained by the mutations discovered here (0.898). In addition, functional redundancy ^75^ and synergistic interaction ^85^ between the paralogus HOX genes could contribute to the additional complexity of the phenotype. However, the genomic region containing the HOXA (OAR04, 68 Mb), HOXC (OAR13, 132 Mb) and HOXD (OAR02, 132 Mb) gene clusters show no signals in the cLDLA analyses without (Model 3) and with (Model 5) candidate mutations as a fixed effect **(Figure 1c** and **d**). Tacking together, to map some remaining causal variants in long-tailed breeds, we will need a design with much higher statistical power than the one carried out in the present study.

Our results provide a comprehensive insight into the genetic variance of tail length in long-tailed Merino sheep and offer information towards direct gene-assisted selection for shorter tails and thus contribute to animal welfare by avoiding tail docking and mulesing in the future. Part of society, which is actively committed to animal welfare, frequently has prejudices against genetic methods in general. Therefore, it should be pointed out once again that this is a natural original genetic variant and that selection in favor of this variant serves to restore the most natural original trait. We would not call it a “repair”, but a “back to the roots”. Additional to commercial and animal welfare aspects this and follow-up study could contribute to a better understanding of embryonal development too. According to Aires, et al. ^86^ quantitative differences within the *Gdf11-Lin28-Hoxb13-Hoxc13* gene network might account for the tail size variability observed among vertebrate species. Thereby, the determination of the tail length could result from the relative intensity or the sequence of the individual network components.

The mechanisms regulating tail size are still not fully understood especially the pathways downstream of *Lin28* and *Hox13* genes in the *Gdf11-Lin28-Hoxb13-Hoxc13* network. Therefore, Aires, et al. ^86^ suggested testing the gene-network parameters in embryos of vertebrate species with different tail sizes. The embryos of a sheep breed with an ancestral short tail and a derived long tail are suitable candidates for studying the control of axis termination. In the meantime, the mapping design could be extended and improved to map genes pointing to further network candidates, possibly downstream mediators of the *HOXB13.*

## Conclusions

From the results of our well-structured mapping population, we concluded that the tail length is a highly heritable trait and depends on many loci with minor effects, whereas variations around *HOXB13* cause the main effect in tail length. This is evident from the only slight reduction in heritability after correcting for this major locus. To detect the additional causal loci a more powerful design is needed. We have shown that variance component analysis in the haplotype-based mixed linear model can be more successful than methods such as GWAS when the number of phenotyped individuals and genotyped markers is not large enough for GWAS. Further, our results indicated that second generation short reeds sequencing technology coupled with the assembly errors can make finding promising structural candidate variants difficult and thus visual examination of the targeted region should always be considered. However, long-read sequencing technology such as PacBio and ONT have already proven to be effective in identifying such SVs. Despite this, we were able to detect an insertion and a SNP within the promotor and exon regions of *HOXB13* as the most plausible but possibly not sole cause for the major effect. Furthermore, our results suggest sheep as model animals for deciphering the mechanism of general interest, e.g. developmental biology and cancer. Finally, we clearly show that selection for shorter tails in economically most important long-tailed Merino breeds is possible without introgression and without negative side effects and that this selection is ethically unproblematic as it leads to an increased frequency of the ancestral and thus for sheep most natural allele.

## Declarations

### Ethics approval and consent to participate

The collection of blood samples for this study was approved by the ethics committee of the Veterinary Faculty of LMU Munich.

All blood samples were taken according to best veterinary practice and under a permit from the Government of Upper Bavaria (permit number: 55.2-1-54-2532.0-47-2016), or from the Regional Council of Gießen, Hassia (KTV number: 19 c 20 15 h 02 Gi 19/1 KTV 22/2020).

### Consent for publication

Not applicable

### Availability of data and materials

The datasets generated during and/or analysed during the current study are available from the corresponding author on reasonable request.

### Authors’ contributions

DL analyzed and interpreted data and drafted the manuscript. EH and KE collected samples and phenotypes and assisted in drafting the manuscript. JK assisted in the interpretation of the analyses. DS and IR coordinated genotyping of the samples and provided data. CM provided expertise about the status and problems of modern sheep breeding and established contacts with breeders. GL provided samples and phenotypes. SK performed capture sequencing and assisted in drafting the manuscript. HB performed and coordinated capture sequencing. MU carried out data analyses and interpretation and assisted in drafting the manuscript. IM conceived and guided this study, provided analysis tools, analysed data and critically revised the manuscript. All authors read and approved the final manuscript.

## Acknowledgements

The authors thank to breeder Hermann Stadler for making his sheep available for measuring and sampling.

